# Quantitative Imaging of the Heterogeneity of Brain Potassium Depletion in Experimental Focal Ischemia

**DOI:** 10.64898/2026.03.13.710182

**Authors:** Alexander Kharlamov, Victor E. Yushmanov, Kirk A. Easley, Boris Yanovski, Stephen C. Jones

**Affiliations:** CerebroScope, the dba entity of SciencePlusPlease LLC, 4165 Blair St., Pittsburgh, PA 15207-1508, USA; Department of Anesthesiology, B’nai Zion Medical Center, Sderot Eliyahu Golomb 47, Haifa, Israel; Previously Department of Anesthesiology, Allegheny-Singer Research Institute, 320 E. North Ave., Pittsburgh, PA, 15212; Department of Biostatistics and Bioinformatics, Rollins School of Public Health, Emory University, 1518 Clifton Road NE, Atlanta, GA, 30322, USA

**Keywords:** acute focal ischemic stroke, potassium imaging, spreading depolarization, peri-infarct depolarizations, brain

## Abstract

**BACKGROUND:** With few exceptions, pathological progression in ischemic stroke is presumed to occur uniformly within the ischemic core region. These exceptions include edema formation, brain tissue [Na^+^] increase, and the qualitative visually observed decrease of brain tissue [K^+^], [K^+^]_br_, all of which occur in peripheral regions of the ischemic core. We hypothesize that [K^+^]_br_ depletion and egress occur heterogeneously in the peripheral compared to the central ischemic core and this heterogeneity is not associated with neuronal degradation.

**METHODS:** Permanent focal ischemia was produced in 13 rats for 2.5-5 h. Brain sections were quantitatively stained for K^+^ to assess variations in [K^+^]_br_ depletion and egress between the peripheral and central ischemic core. Reflective change and microtubule-associated protein 2 (MAP2) stained sections were used to identify the ischemic region and relate neuronal pathology to [K^+^]_br_ variations.

**RESULTS:** The mean value of normal cortex [K^+^]_br_ was 96 mEq/kg and of K^+^-egress in all ischemic regions over time was 12.2 mEq/kg/h, consistent with measurements from other studies. Significant differences in exaggerated K^+^-depletion (p<0.001) and egress (p=0.010) occurred in 56% of the peripheral compared to central ischemic core regions suggesting accelerated K^+^-egress from 0 to 2.5 h. Unlike [K^+^]_br_, there was no difference between the MAP2 immunoreactivity in K^+^-depleted and non-K^+^-depleted peripheral ischemic core regions (p=0.83, p=0.16, respectively).

**CONCLUSIONS:** While confirming previous results of quantitative losses of [K^+^]_br_ in the ischemic core, we additionally show using quantitative imaging that K^+^ dynamics within and between the peripheral and the central ischemic core are heterogeneous and not related to MAP2-assessed neuronal structural integrity. Insufficient K^+^ in K^+^-depleted peripheral ischemic core regions might limit spreading depolarization-mediated infarct expansion and not allow restoration of the parenchymal membrane potential even if the functionality of the Na^+^,K^+^-ATPase is restored. Further study of differing K^+^-dynamics within the ischemic core might lead to a better understanding of ischemic stroke pathophysiology.

## Introduction

Sodium and K^+^ dynamics in ischemic stroke have been viewed as important issues because they are the major cations that maintain membrane potential via the Na^+^,K^+^-ATPase and whose transient intra-intercellular disequilibrium subserve ischemic depolarization. Many experimental studies have used increasingly more sophisticated methods for their determination^1^, including atomic absorption spectroscopy^2,3^, ion-sensitive electrodes^4^, non-hydrogen-based MRI^5–8^, ion-sensitive fluorochrome imaging^9^, and genetically encoded fluorescent probes^10^. Although Na^+^ and K^+^ can be viewed as complementary cations, with K^+^ being intracellular and Na^+^ extracellular, the vast majority of experimental effort has been directed at Na^+^ dynamics. Na^+^ accumulation and subsequent edema is considered one of the principal mechanisms of ischemic stroke pathophysiology^11,12^ and is the major pathological factor of brain-edema instigated tentorial herniation^13,14^, the most egregious event in the progression of ischemic pathology. This focus on Na^+^ has left the consideration of K^+^ dynamics in the background. However, although Na^+^ is important in the immediate pathological consequences of ischemic stroke, K^+^ is equally important if recovery issues are to be considered. Because K^+^ moves from intra- to extra-cellular during depolarization, it is exposed to the brain’s exit systems, in contrast to Na^+^’s cellular entrapment. Thus, post-ischemic measurements of K^+^ (of course almost always made in concert with Na^+^) show a consistent and excessive loss of K^+^ from the brain^15–24^ as presented in Table 1.

**Table 1.**
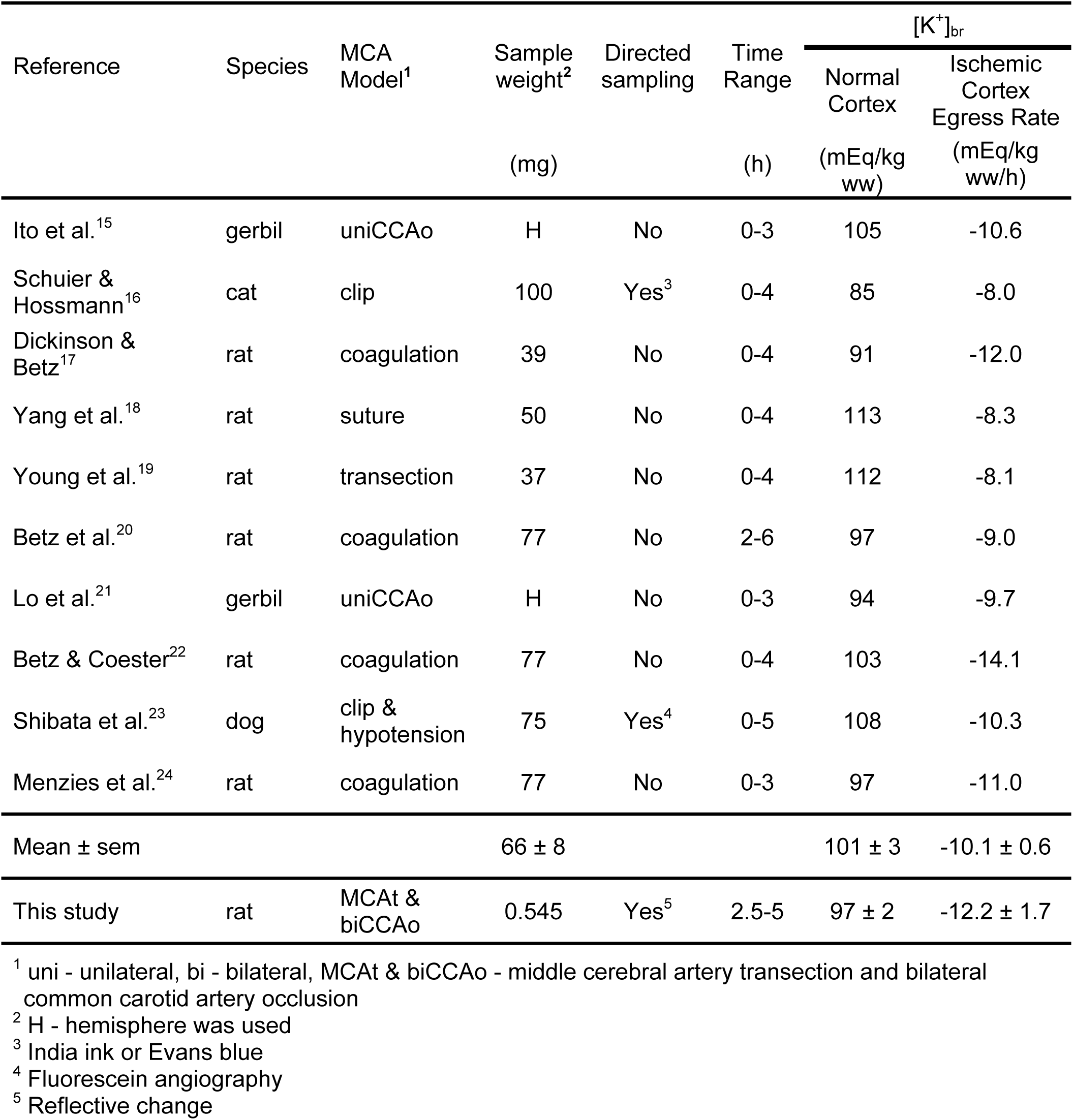
Comparison with other measurements of [K^+^]_br_ normal cortex and ischemic cortex egress rate in models of focal ischemia.

These measurements of K^+^ in acute ischemic stroke^15–24^ were made with the assumption that the ischemic core [K^+^]_br_ was homogeneous. However, there have been reports of intra-core heterogeneity in ischemic stroke. These include observations in the peripheral regions of ischemic core of increased edema, lowered brain tissue [K^+^], [K^+^]_br_, and increased [Na^+^]_br_^25^, a maximal rate of [Na^+^]_br_ increase^7,26^, transient blood-brain barrier (BBB) breakdown to gadolinium diethylenetriaminepentaacetic acid (Gd-DTPA)^27^, decreased reflective change^28^, decreased MAP2 immunoreactivity (IMR)^28–30^, and decreasing [K^+^]_ex_^31^. These measurements suggest that the peripheral ischemic core has a special and unique significance during this acute period that relates to the cortical spreading depolarizations (SDs) that occur repetitively around the edge of the ischemic core^32,33^.

To expand our understanding of K^+^ dynamics to include a more intra-ischemic core view, we propose to use quantitative K^+^-imaging^34^. Previous methods used for the measurements in Table 1 cannot address the regionality within the ischemic region needed to assemble a complete picture of K^+^-related ischemic core pathology. However, quantitative K^+^-imaging^34^ permits the examination of our hypotheses concerning the regional and temporal differences in the clearance of K^+^ from ischemic tissue: 1) That at early times after ischemia, K^+^ egress from the peripheral and central ischemic core are different; and 2) That these differences in K^+^ dynamics are not based on neuronal integrity assessed by calpain-mediated microtubule disassembly via MAP2 IMR changes.

Answers to these hypotheses could help us understand and mitigate K^+^ related stroke issues. As SDs continue to occur in the initial several hours after stroke onset^35^, more K^+^ becomes extracellular, and more and more K^+^ is lost from the brain. There are at least two consequences of this K^+^ loss that are equally important when compared to Na^+^ accumulation: K^+^ dependent 1) neuronal recovery; and 2) SD ignition and propagation. Even if energy supply can be resupplied to marginally ischemic tissue by the reestablishment of Na^+^,K^+^-ATPase activity, without K^+^, the membrane potential cannot be reestablished. If appropriate [K^+^]_in_ levels cannot be restored to allow SD’s depolarization to occur, even though K^+^ is not a primary factor in SD initiation^36,37^, then eventually SDs will become less likely.

## Methods

### Animal procedures

Animal procedures were performed in compliance with the National Institutes of Health “Guide for the Care and Use of Laboratory Animals” and were approved by the Institutional Animal Care and Use Committee, DHS/OLAW Assurance #A3693-01. Thirteen Sprague/Dawley male rats (285 ± 7 g, mean ± sem) were anesthetized with isoflurane (2.5% induction and 1.0 ± 0.5% maintenance), 30% oxygen and balance nitrous oxide, intubated, and artificially ventilated (Harvard Apparatus, model 683, South Natick, MA). During maintenance anesthesia, the concentration of isoflurane was titrated by monitoring the arterial blood pressure response to tail-pinch. Body temperature was maintained at 37°C by a rectal probe and servo-controlled heating lamp (YSI 74, Yellow Springs, OH). Venous and arterial femoral catheters (PE50) were inserted. Arterial blood gases (PaCO_2_, PaO_2_), and pH were determined with a portable blood analyzer (i-STAT, Heska, Loveland, CO). Arterial blood pressure was continuously monitored. Pancuronium (Gensia Sicor Pharmaceuticals, Irvine, CA) was administered hourly (0.7 mg/kg, IM). Physiological variables and laser Doppler flowmetry (LDF)-determined cerebral blood flow (CBF) were digitally recorded with Windaq software (Dataq, Inc., Columbus OH).

### Model of brain ischemia

The animals were subjected to permanent focal ischemia produced by right middle cerebral artery transection (MCAt) midway between the olfactory tract and inferior cerebral vein, followed by permanent bilateral common carotid artery occlusion (biCCAo)^26,38^ or, in one animal, MCA permanent intraluminal suture occlusion^39^. CBF was monitored by LDF (λ = 780 nm, LaserFlo BPM403A, Vasamedics, Inc., St. Paul, MN) with the probe firmly placed in a 1-mm diameter skull bone window over the right MCA territory -4 mm from bregma and 5 mm from the midline (see Fig. 1A). CBF was expressed as the percent of pre-ischemic control after correction for biological zero obtained at the end of the experiment^40^. CBF was not recorded in two animals due to technical issues. Animals were killed at 15 minute intervals between 2.5 and 5 h after stroke onset (two each at 2.75 and 4.5 h, and one at each of the other 9 times). The brains were removed and frozen over dry ice and stored at -70°C. In the MCA suture animal, the whole head was frozen and the brain chipped out.

**Figure 1.**
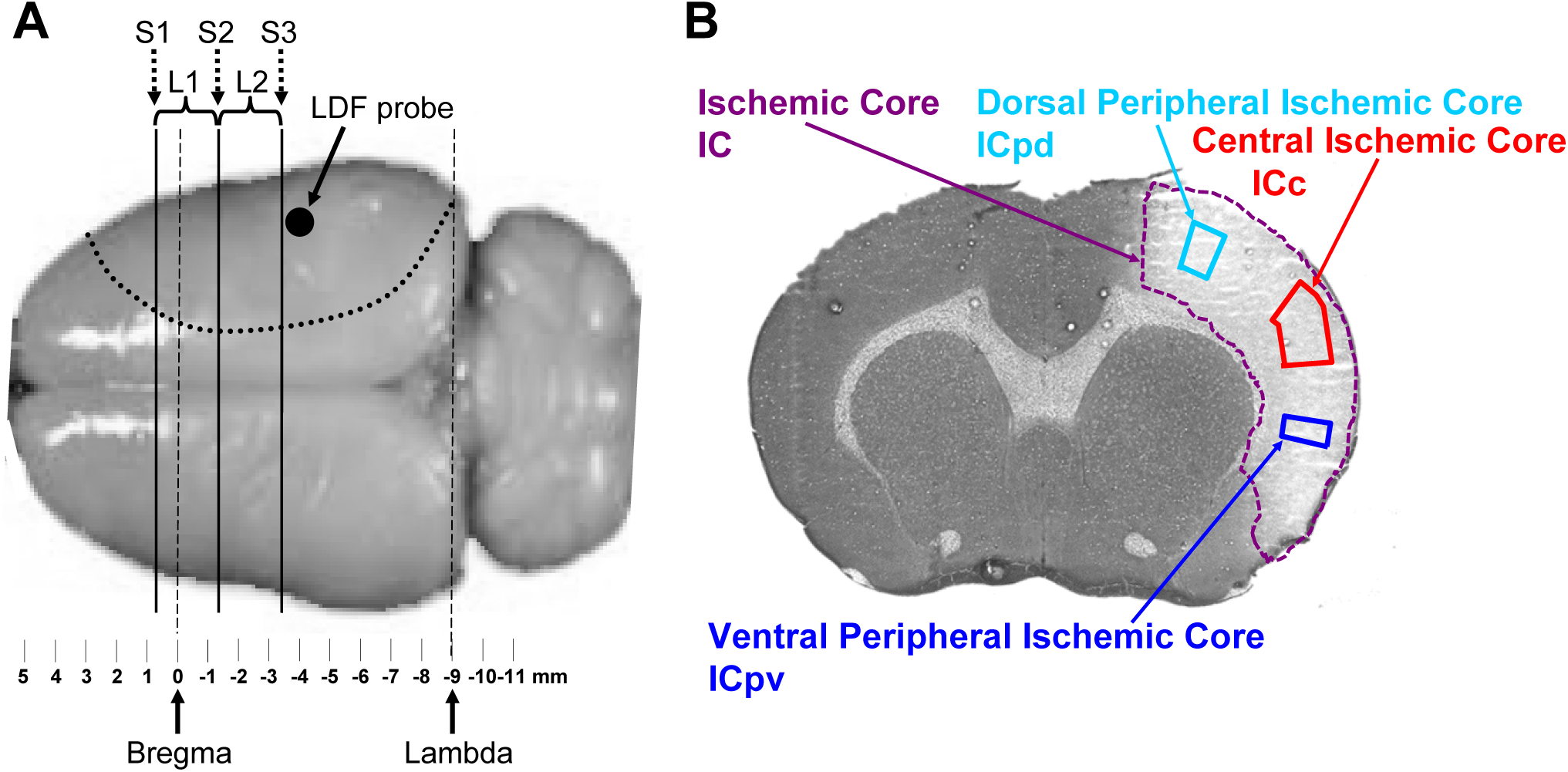
**(A)** A dorsal view of a typical rat brain with bregma and lambda showing: 1) the selection of the 2-mm-thick slices for the first and second series of micropunch samples at two levels, L1 and L2; 2) the three positions, S1, S2, and S3, for the 35-µm sections taken for K^+^-staining and cortical ribbon analysis, MAP2 IMR, and reflective change; and 3) the laser Doppler flowmeter (LDF) probe position. The dashed line indicates the approximate range of the middle cerebral artery distribution that typifies the infarct area. **(B)** Histo-K^+^-stained coronal section showing the ischemic core (IC, violet), as confirmed by reflective change and MAP2 IMR in adjacent sections, and its division into regions-of-interest for dorsal peripheral (ICpd, light blue), ventral peripheral (ICpv, blue), and central ischemic core (ICc, red) regions, as identified by variations in [K^+^]_br_. The ischemic penumbra would be located just outside the dorsal and ventral peripheral ischemic core in the regions where [K^+^]_br_ is approaching normal, non-ischemic levels (see Fig. 2H).

### Brain processing

Brain tissue samples were collected in the microtome-cryostat (Minotome, IEC, Needham Heights, MA) at -7°C using a micropuncher (2 x 0.5 mm ID) for flame photometry analysis of [K^+^] ^34,41^. Micropunch samples (depth ∼2 mm, 0.545 ± 0.005 mg, 14 punches/level x 2 levels/animal x 13 animals) from ischemic and contralateral (normal) brain areas were taken at two different rostro-caudal levels as shown in Fig. 1A: L1 (+0.7 to -1.3 mm from bregma) and L2 (-1.3 to -3.3 mm from bregma). The K^+^-content of brain tissue samples was determined by emission flame photometry at 766 nm (IL943 flame photometer, Instrumentation Laboratory, Lexington, MA) and expressed as K^+^ brain concentration, [K^+^]_br_, in mEq/kg^34^. The change of surface reflectivity, or reflective change^28^, on adjacent dried coronal sections (35 µm, Fig. 2A) provided unequivocal differentiation of ischemic from normal cortex during micropunch sampling and confirmed the extent of focal ischemic damage. The images of the cut-face of the brain (Fig. 2B) at each level (S1, S2, and S3, Fig. 1A) were digitized for later alignment of the micropunch locations with adjacent histo-K^+^-stained sections (Fig. 2C). Additional adjacent sections were immunoreactively stained for the cytoskeletal protein MAP2^28^, as shown in Fig. 2G.

**Figure 2.**
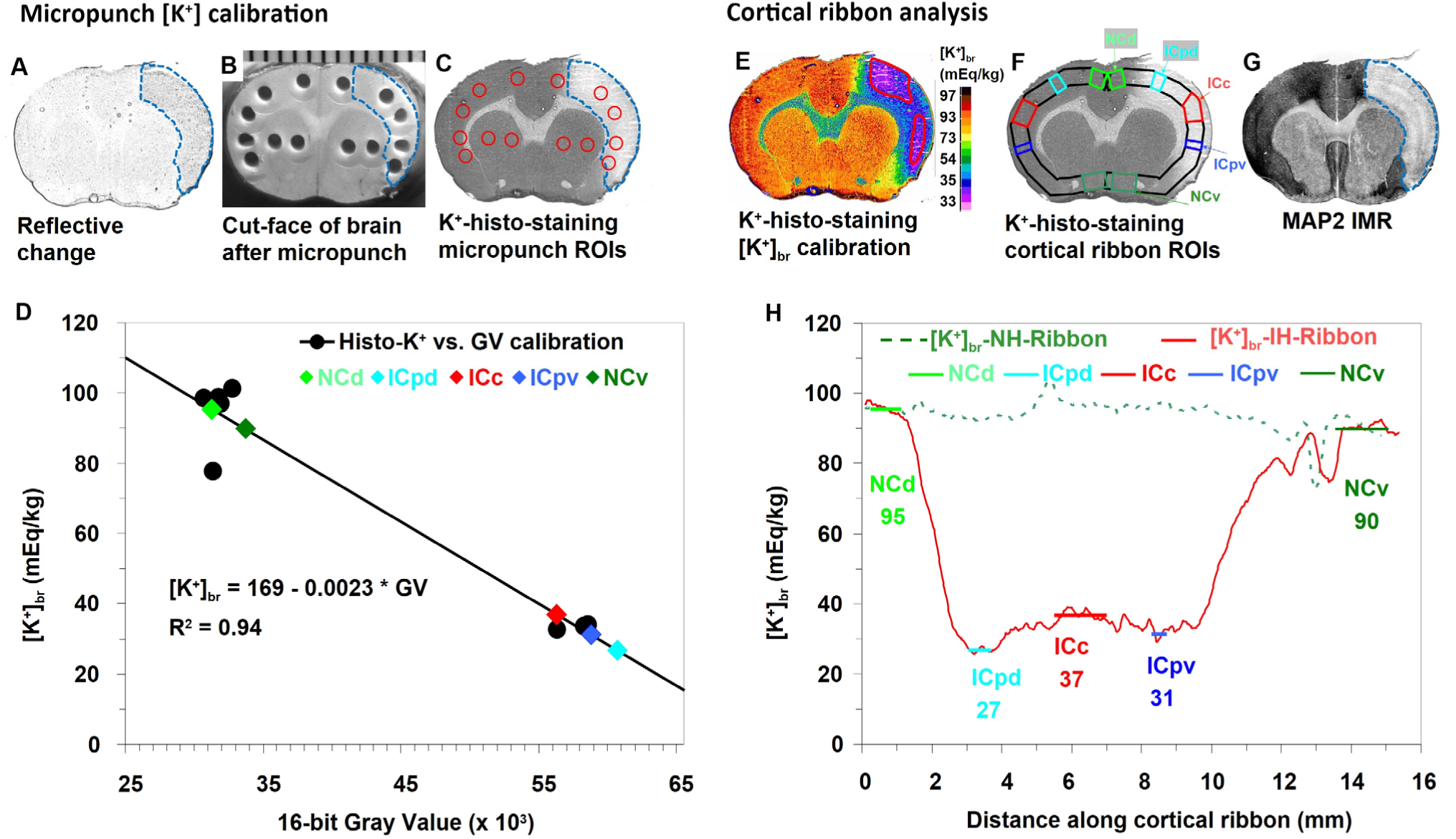
The sequence of brain processing steps and other procedures for adjacent sections and images at the S1 level (see Figure 1). All images are aligned and processed digitally. In panels **A**, **B**, **C**, and **G** the blue dashed line indicates the infarct area. **(A)** The reflective change in a coronal brain section used to guide the micropunch locations to normal cortex, ischemic cortex, and subcortical regions. **(B)** The cut-face of the brain in the cryostat showing the micropunch hole positions used for ROI placement in panel **C**. The mm scale placed at the brain section level is at the top of the Figure. **(C)** Histo-K^+^-section showing the micropunch ROIs (red circles) obtained from panel **B** that are used to create the calibration (see panel **D**) of the gray value (GV) by the flame photometry-determined [K^+^]_br_ in each micropunch. **(D)** The calibration curve from the linear regression of micropunch ROI GVs vs. flame photometry [K^+^]_br_ values (circles) was used to convert the GVs of the histochemical image to [K^+^]_br_. The [K^+^]_br_ values obtained from the GVs of the cortical ribbon ROIs (panels **F** and **H**) are shown along the curve as colored diamond symbols. **(E)** Pseudo-color coded histo-K^+^-image based on the calibration curve shown in panel **D** with the color representing [K^+^]_br_ in mEq/kg as shown on the calibration bar. The red line in the ischemic right cortical region indicates the 36 mEq/kg isocontour which is the K^+^-depletion breakpoint defined in Figure 3. **(F)** The histo-K^+^ image shows the cortical ribbon for the normal and ischemic hemispheres (black lines) with the ROIs for dorsal normal cortex (NCd, light green), dorsal peripheral ischemic core (ICpd, light blue), central ischemic core (ICc, red), peripheral ventral ischemic core (ICpv, dark blue), and ventral normal cortex (NCv, dark green). **(G)** Section stained for MAP2 IMR to confirm the extent of the ischemic lesion and to assess possible MAP2 IMR decrements in the peripheral ischemic core for comparison to K^+^-depletion. **(H)** The [K^+^]_br_ profiles along the cortical ribbon for the ischemic (IH-ribbon, orange curve) and non-ischemic hemispheres (NH-ribbon, dotted-green curve). The horizontal lines indicate the position of the five ROIs (**F**, described above) along the ischemic hemisphere cortical ribbon and their [K^+^]_br_ values. The dorsal (ICpd) and ventral (ICpv) peripheral ischemic core regions, located at the edges of the ICc, demonstrated larger decreases of [K^+^]_br_ than the decrease in the ICc. In this animal, the ICpd and ICpv regions were 72% and 84% of the ICc value, respectively, so the ICpd region, but not the ICpv, was classified as K^+^-depleted because it was <78%, the K^+^-depletion breakpoint, as described in the text and shown in Figure 3.

### Histological K^+^-staining and [K^+^]_br_ calibration

The quantitative histochemical method for brain K^+^-staining described previously^34^ was used. This method is based on the chemical reaction of K^+^ with sodium cobalt hexanitrite, Na_3_Co_2_(NO)_6_, with formation of potassium cobaltinitrite crystals. The barely visible yellow crystalline precipitates of potassium cobaltinitrite transform into easily detectable black cobalt sulfide crystals by adding ammonium sulfide in a cold bath kept below 4°C^34,42,43^. The sections histochemically stained for K^+^ (histo-K^+^-stained) were digitized and calibrated using flame photometry-determined [K^+^]_br_ values from micropunch samples (Fig. 2D). The optical densities of each histo-K^+^-stained section were separately calibrated against flame photometry determined [K^+^]_br_ from micropunch samples taken directly from the cut-face left after that microtome section was taken.

### Time-course analysis of cortical-ribbon K^+^-depletion

In the regions of reflective change and depressed MAP2 IMR, [K^+^]_br_ dynamics were analyzed in the acute period after experimental ischemic stroke in three coronal sections (S1, S2, and S3 in Fig. 1A). The calibrated histo-K^+^ images (Fig. 2E) were analyzed for spatial changes of [K^+^]_br_ using the ribbon tool of the MCID imaging system (Version 7, InterFocus Imaging Ltd., Linton, United Kingdom). The ribbon tool produced a plot of the average [K^+^]_br_ within the cortical rim from the dorsal hemispheric fissure to the ventral midline and allows the placement of within-ribbon regions-of-interest (ROIs). The width of the ribbon was defined by the plateau of the GV profile of the density across the cortex layers indicating uniform staining. Using this width of 0.9 mm, the cortical profiles of [K^+^]_br_ along the ischemic and non-ischemic hemispheres were obtained (see Fig. 2F and 2H).

The 39 [K^+^]_br_ ischemic hemisphere-ribbons (3 sections per animal x 13 animals) showed two distinct patterns of their [K^+^]_br_ cortical profiles. One pattern (25 ribbons) had a wide ischemic plateau often with low [K^+^]_br_ at one or both peripheral ischemic core regions. The other pattern (14 ribbons) had only a sharp minimum without a wide central ischemic plateau region so a central ischemic core (ICc) region could not be identified. The [K^+^]_br_ profile of these patterns, which appeared as a “wedge”, occurred because the cortical ribbon “cut” was positioned at the caudal or rostral edges of the ischemic region. The [K^+^]_br_ in these cortical ribbons with “wedge” patterns were therefore excluded from analysis because K^+^ loss at the peripheral parts of the ischemic core could not be compared to that in the central ischemic core.

Differences in [K^+^]_br_ between the ICc and the dorsal and ventral peripheral ischemic core (ICpd and ICpv) regions (Fig. 2F) were apparent from assessment of the [K^+^]_br_ ischemic hemisphere-ribbons of the histo-K^+^-stained sections: [K^+^]_br_ was depressed in some but not all peripheral ischemic core regions. Further [K^+^]_br_ analysis was driven by this unusual pattern of excessively depressed [K^+^]_br_, termed K^+^-depletion, in the peripheral ischemic core in relation to the central ischemic core.

In the remaining 25 cortical ribbons (out of the original 39), five different within-ribbon ROIs were selected in the ischemic hemisphere-ribbon: a) two normal cortex regions of parasagittal and pyriform parts of the ischemic hemisphere (dorsal normal cortex, NCd, and ventral normal cortex, NCv, respectively); b) the ICc; and c) two ROIs at the periphery of the ischemic core, one dorsal (ICpd) and one ventral (ICpv) region identified by a consistent depression of [K^+^]_br_ or at those same positions in sections without such consistent depression (see Fig. 2F and 2H). Peripheral ischemic core regions (ICp) were selected based on their [K^+^]_br_ distribution according to our hypotheses as there is no known morphological attribute or criteria that provides such guidance. The [K^+^]_br_’s of the ICpd and ICpv regions were combined into one group, ICp, for additional analysis, as they were deemed statistically similar (p>0.59, paired t-test). Five corresponding homotopic regions were chosen in the non-ischemic hemisphere ribbon.

To analyze K^+^-egress from these ICp regions, their [K^+^]_br_, [K^+^]ICp, was expressed as a percentage of that in the ICc region, ([K^+^]ICc), based on the assumption that the [K^+^]_br_ of these peripheral ischemic core regions was originally equal to that of the central ischemic core:

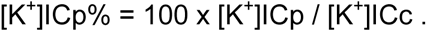

Histogram analysis was then used to explore variations in K^+^-depletion within the ischemic core. The frequency distribution of [K^+^]ICp% (n = 50) revealed an apparently bimodal distribution. To assess this distribution for bimodality, two models were compared; a bimodal model with two Lorentz curves and another with just one Lorentz curve (see Fig. 3). The trough of the bimodal frequency distribution, the [K^+^]ICp% value at the intersection of the two Lorentz curves, was used to establish a breakpoint between peripheral ischemic core regions with (ICp-DP) and without (ICp-ND) K^+^-depletion.

**Figure 3.**
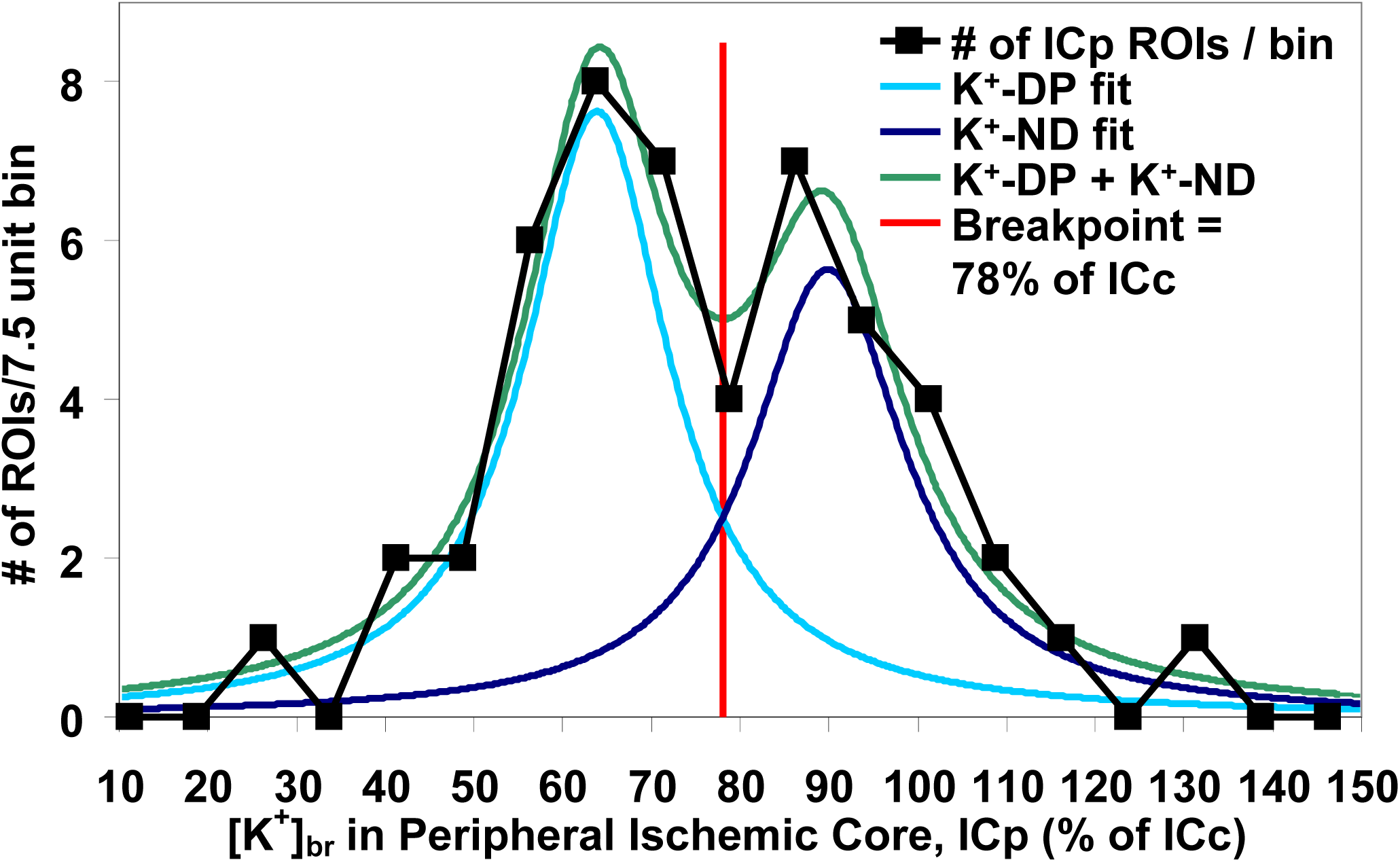
Histogram analyses of [K^+^]_br_ in the combined peripheral ischemic core (ICpd and ICpv) and central ischemic core (ICc) cortical-ribbon ROIs. Frequency analysis of % changes of [K^+^]_br_ in the combined peripheral ischemic core regions (black squares and line, n = 50) revealed a bimodal distribution with medians of 64% (light blue line) and 90% (dark blue line). The minimum between the two peaks of the summed distribution (green line) was chosen as the breakpoint (78%, vertical red line) to distinguish the K^+^-depleted peripheral ischemic core regions from those that retained [K^+^]_br_ similar to the ICc values.

Finally, [K^+^]_br_ values of the NC, ICc, ICp-DP and ICp-ND regions were plotted vs. time after stroke onset (Fig. 4). Each of these curves was assessed for linearity and differences between their slopes and intercepts were statistically tested. Forty-four peripheral ischemic core regions were available for analysis: 6 regions from suture model animal (2 peripheral ischemic core regions * 3 sections) were excluded from the 50 peripheral ischemic core regions. This animal was excluded from the time course analysis because the suture MCA model used in this animal has a slower progression of cortical ischemic pathology^44^ in comparison to the direct distal occlusion of the MCA^26^ used in all other animals.

**Figure 4.**
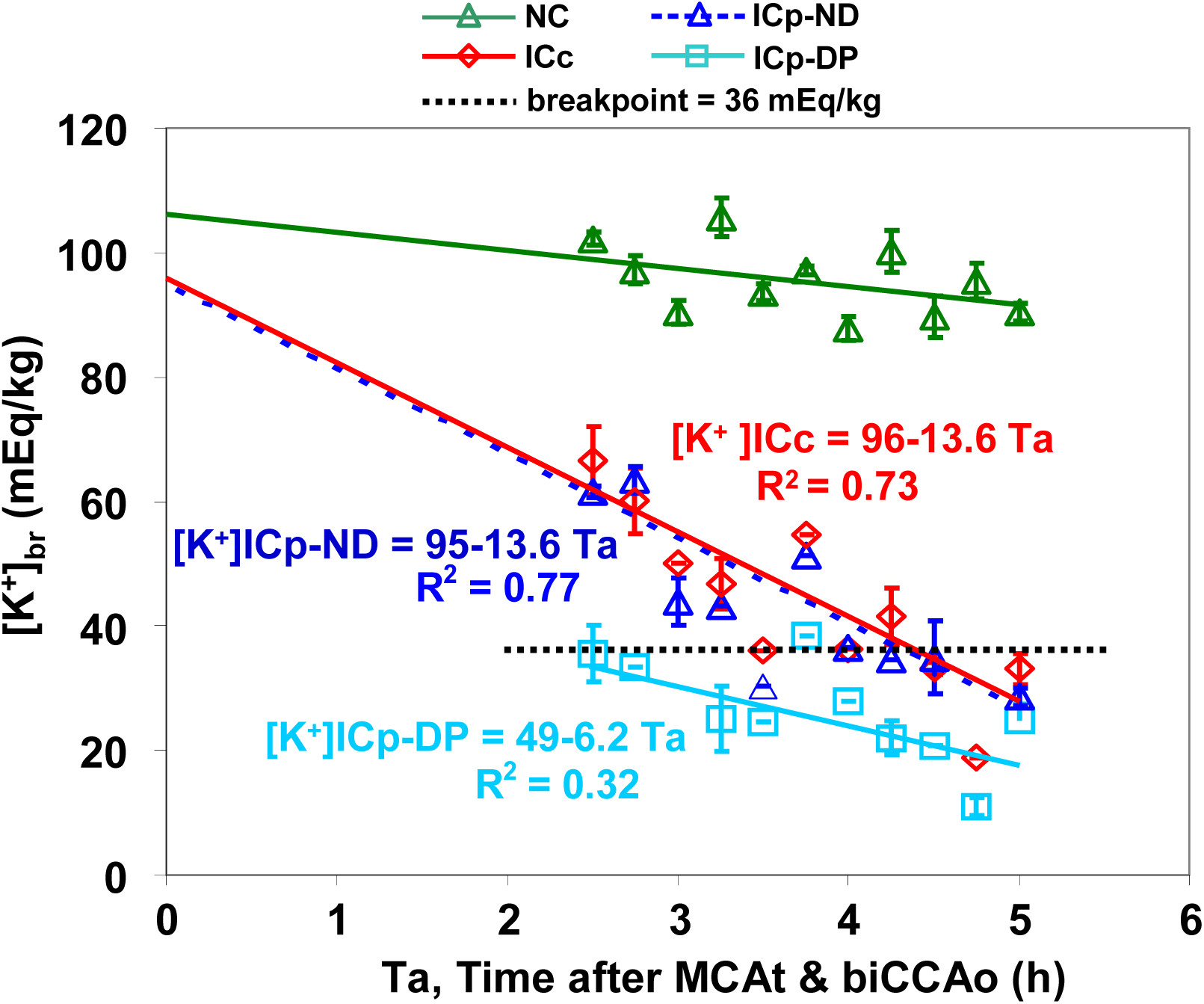
Time course of [K^+^]_br_ (mean ± sem) from the cortical ribbon ROIs in normal cortex (NC), central ischemic core (ICc) and peripheral ischemic core (ICp) regions between 2.5 and 5 h with their linear regression curves. The ICpd and ICpv peripheral ischemic core regions have been combined, but then divided into K^+^-depleted (ICp-DP) and non-K^+^-depleted (ICp-ND) groups on the basis of their [K^+^] expressed as a percent of the ICc value. This dividing line, or breakpoint, is shown converted to mEq/kg as the horizontal dashed line at 36 mEq/kg. The time course of the combined NCd and NCv regions (NC, green line) shows stability of [K^+^]_br_, whereas the ICc (red line), ICp-DP (light blue line), and ICp-ND (dark blue dotted line) time courses all show a significant decrease over time, with the slopes of the ICp-ND regions and the ICc being similar (p =0.99). The NC, ICc, and ICp-ND curves have been extrapolated to the stroke onset time, Ta = 0. The slope of the ICp-DP region is significantly different from that of the ICc (p = 0.010) and the ICp-ND (p = 0.0071) curves. This lower slope of the ICp-DP curve 2.5 h after Ta is due to the accelerated K^+^-egress from Ta = 0 to 2.5 h, as its value at Ta = 0 must be ∼100 mEq/kg.

### Immunostaining for cytoskeletal protein, microtubule-associated protein-2 (MAP2)

The decrease of IMR for the cytoskeletal protein MAP2 was used as a marker of early ischemic damage^28–30^. The MAP2 IMR was detected with anti-MAP2 primary antibody (HM2, 1:500, Sigma-Aldrich, St. Louis, MO) using our previously published procedure^28^ on adjacent sections to those histo-K^+^-stained. MAP2 immunostaining in frozen sections can lead to artifacts, so only 27 immuno-stained sections (out of 39 = 3 sections per animal x 13 animals) were technically acceptable for analysis (two animals had none, six animals had 2, and the remaining 5 had 3 each that were acceptable). MAP2 IMR was normalized by expressing the ratio of the gray value (GV) in each ROI to the average GV of normal cortex for each section to correct for variations in section thickness, tissue processing, and staining intensity. Decrements of MAP2 IMR and [K^+^]_br_ were compared using the transposed cortical ribbon ROIs of the histo-K^+^-stained sections (Fig. 2F) onto the MAP2 IMR sections (Fig. 2G) as presented in Fig. 5.

**Figure 5.**
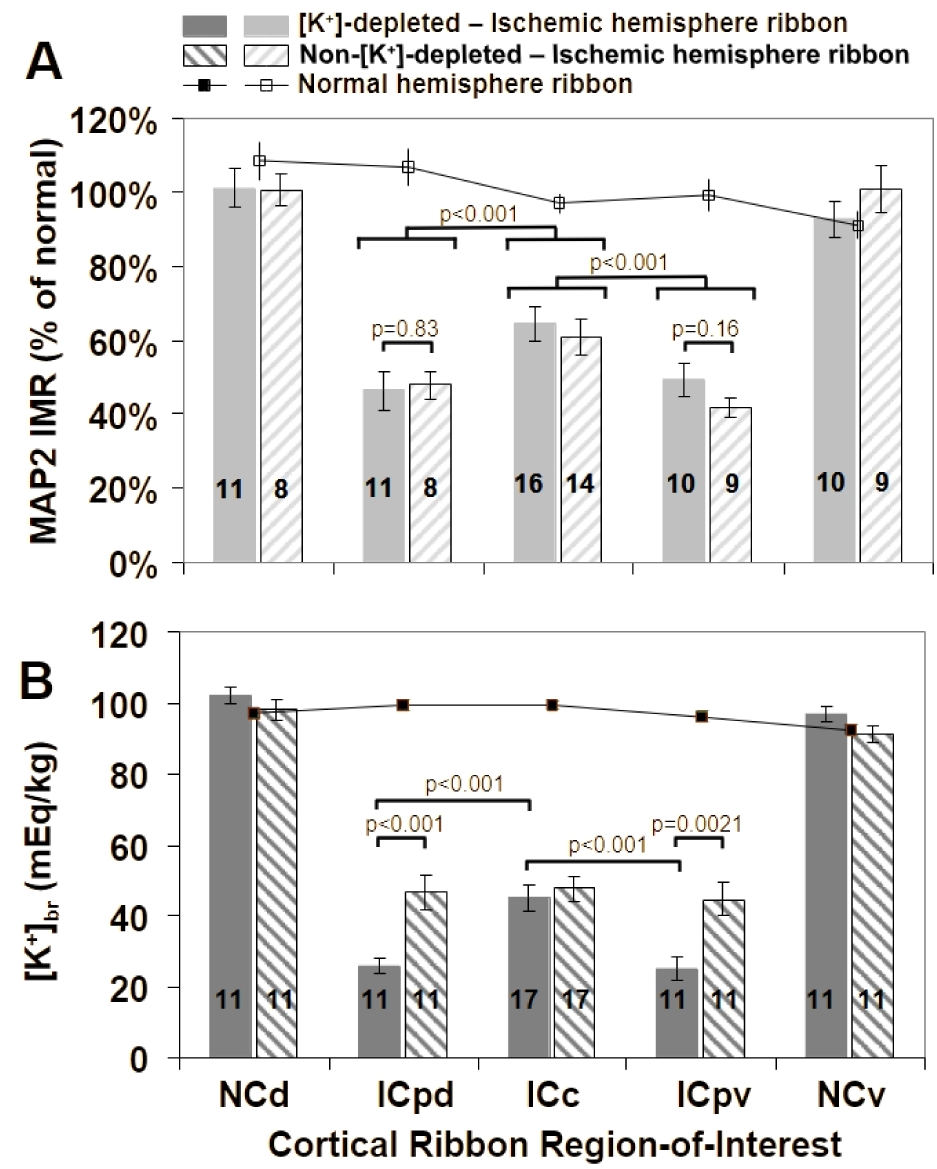
Analysis of [K^+^]_br_ and MAP2 IMR from ROIs in the normal and ischemic hemispheric cortical ribbons for the purpose of testing whether the ICpd and ICpv regions differ in their K^+^-depletion and MAP2 IMR. The normal hemisphere values (mean ± sem) are plotted as the solid lines with filled squares. The ischemic cortical ribbon ROIs have been divided into those whose ICpd and ICpv ROIs are K^+^-depleted (solid bars with n’s shown inside each bar) and non-K^+^-depleted (stripped bars) for both panel **A**, the MAP2 normalized IMR values and panel **B**, showing the [K^+^]_br_ values. The normal cortex values are divided based on whether their associated peripheral ischemic cortex ROI values are K^+^-depleted or non-K^+^-depleted. **(A)** The MAP2 IMR normalized to the gray value in normal cortex shows no difference between the ICpd and ICpv K^+^-depleted and non-K^+^-depleted values (unpaired t-test, p = 0.83, p = 0.16, respectively), but a significant decrease in ICpd and ICpv vs. ICc IMR (paired t-test, p < 0.001 for both). **(B)** A significant [K^+^]_br_ decrease in the K^+^-depleted peripheral ischemic core regions (ICpd and ICpv) was observed in comparison to: (1) the [K^+^]_br_ in the central ischemic core (ICc) (two-tailed paired t-test, p < 0.001 for both); and (2) the non-K^+^-depleted regions (unpaired two-tailed t-test, p < 0.001, p = 0.0021, respectively). However, there was no significant difference between the non-K^+^-depleted and ICc values.

### Statistics

Two-tailed paired and unpaired t-tests were used to test differences between CBF over time and between the ICc and ICp ROI values of [K^+^]_br_ and normalized MAP2 IMR. The distribution of [K^+^]ICp% values was tested for bimodality by comparing the fitted Lorenz distribution of all the data to the sum of the bimodal Lorenz distributions using an F-test and a second-order Akaike Information Criterion (AIC) test using OriginPro (OriginLab Corporation, Northampton, MA). Differences between slopes of the linear regression analysis of the [K^+^]_br_ time-courses were evaluated from the t-statistic calculated by their difference divided by the RMS of their standard errors. These linear regression analyses were performed on pooled ROI data, which assumes independence between samples (ROIs are nested within animals). Consequently, the between-animal variance may not be negligible, and the standard errors used for slope comparisons via the t-statistic may be underestimated with the caveat that precise estimates of the two variance components would be limited with only 13 animals. Values are presented as mean ± sem. Significance was assumed for p < 0.05.

## Results

Physiological variables (mean arterial blood pressure, blood pH, PaCO_2_ and PaO_2_) were within normal limits before ischemia and were stable after ischemia (Table 2). After ischemia, the CBF over the cortical area of the right MCA territory decreased to 16% ± 3% (mean ± sem, n = 11) of control (pre-ischemic) values (Table 2). At the end of the experiment, the mean CBF was 28% ± 7%, showing an insignificant trend higher than the immediate post-ischemic value (p = 0.063, Table 2), consistent with potentially increased collateral circulation in 3 of the animals.

**Table 2.** Physiological variables at various experimental stages.

| **Table 2** Physiological variables at various experimental stages |  |  |  |  |  |
| --- | --- | --- | --- | --- | --- |
| Time | MABP<br>(mmHg) | pH<br>(-) | PaCO<sub>2</sub><br>(mmHg) | PaO<sub>2</sub><br>(mmHg) | CBF<br>(%) |
| Before ischemia | 108±2 | 7.422±0.009 | 36±1 | 98±3 | 100 |
| ~3 min after ischemia | 115±2 | 7.391±0.016 | 36±1 | 95±4 | 16±3 |
| 120 min after ischemia | 104±3 | 7.413±0.017 | 35±1 | 97±8 | 25±7 |
| End of experiment | 104±2 | 7.400±0.017 | 34±1 | 101±9 | 28±7* |

MABP, mean arterial blood pressure; CBF, cerebral blood flow
\*, p = 0.063 from the immediate post occlusion value at ~3 min (n = 11)
Mean ± sem, n = 11 to 13

Ischemic damage from reflective change (Fig. 2A) and MAP2 IMR images (Fig. 2G) was observed only in the right cortex within the MCA distribution, extending from the frontoparietal cortex to the parts of the parasagittal and pyriform cortices. The normal (non-ischemic) cortical and subcortical structures on histo-K^+^-sections were uniformly stained a dark brown color with white matter less intensely stained than gray matter. The ischemic cortex demonstrated a distinct loss of histo-K^+^-staining. However, the peripheral ischemic core regions often, but not always, showed more intense loss of K^+^ staining, termed K^+^-depletion (Fig. 2E and 2H).

### Histogram analysis for K^+^-depletion

Histogram analysis of the [K^+^]ICp% values was used to compare the lower [K^+^]_br_ values in the dorsal and ventral peripheral ischemic core regions (the K^+^-depleted group, termed ICp-DP, to the non-K^+^-depleted regions, termed ICp-ND). The [K^+^]ICp% values conformed to a bimodal distribution (Fig. 3, p < 0.001, F-test), with an AIC weight ratio indicating that the bimodal fit is 7.5 x 10^7^ times more likely to be correct than a single function fit. A breakpoint [K^+^]ICp% value of 78% separated the ICp regions into two groups: those below, K^+^-depleted (ICp-DP), and those above, non-K^+^-depleted (ICp-ND). In this separation of peripheral ischemic core regions with and without K^+^-depletion, the right-hand peak of Fig. 3 represents [K^+^]_br_’s with values similar to that of the ICc regions (median [K^+^]ICp% of 90%), and the other, with a median [K^+^]ICp% of 64%, represents ICp regions that have undergone K^+^-depletion.

The breakpoint of 78% (Fig. 3, vertical red line) corresponds to 36 mEq/kg (calculated as 0.78 of the mean [K^+^]ICc, 45.8 ± 3.1 mEq/kg). K^+^-depletion was noted in all but 2 of the 13 animals and in 28 (or 56%) of the 50 ICp regions in the 25 cortical ribbons available from both the MCAt&biCCAo and MCA-suture models.

The comparison of [K^+^]_br_ in the peripheral ischemic core regions with and without K^+^-depletion clearly indicates that the [K^+^]_br_ of the K^+^-depleted ICpd and ICpv regions are approximately one half (25.9 ± 2.3 mEq/kg, n = 11 and 24.9 ± 3.4 mEq/kg, n = 11) of the ICc value (45.0 ± 3.5 mEq/kg, n = 17, p < 0.001 for both, see Figure 5B). Most importantly, the [K^+^]_br_ in the dorsal and ventral peripheral ischemic core regions without K^+^-depletion (46.7 ± 4.8 mEq/kg, n = 11, p = 0.86 and 44.6 ± 4.8 mEq/kg, n = 11, p = 0.59, unpaired t-tests) are approximately equal to the ICc value (47.7 ± 3.6 mEq/kg, n = 17), bolstering our hypothesis that no accelerated K^+^-egress occurred in these particular peripheral ischemic core regions.

### K^+^-egress - time course

For comparison to others’ results, none of which address regional variations within the ischemic core, all the ischemic cortex [K^+^]_br_ values (ICpd, ICpv, and ICc) were combined into one group, [K^+^]IC. The linear regression curve of these combined ischemic core regions over the time from stroke onset, Ta, was [K^+^]IC = 83.8 - 12.2 * Ta, with a K^+^-egress rate of 12.2 mEq/kg/h, as presented in Table 1.

The time courses of K^+^-egress from 2.5 to 5 h are presented in Fig. 4. The stability of [K^+^]_br_ in normal cortex and the varying egress rates of K^+^ from the other regions are clearly shown for normal cortex, non-K^+^-depleted peripheral ischemic core, K^+^-depleted peripheral ischemic core, and central ischemic core. The slope of the K^+^-depleted curve differed significantly (p = 0.0071) from that of the non-K^+^-depleted curve and the ICc curve (p = 0.010). Quantitatively, K^+^ was lost at the same linear rate in the ICc regions as in the non-K^+^-depleted peripheral ischemic core, ICp-ND, regions, with a non-significant difference between their slopes (p = 0.99). When taken together, these results indicate that accelerated K^+^-depletion occurred in the ICp-DP regions during the first 2.5 h after stroke onset and then became slower compared to the ICc and ICp-ND regions.

### K^+^-depletion - relation to MAP2 IMR

The [K^+^]_br_ cortical ribbons and ROIs were transposed to the aligned MAP2 images and used to assess whether the loss of normalized MAP2 IMR was associated with the loss of K^+^. MAP2 IMR was decreased in the peripheral ischemic core regions compared to their corresponding NCd and NCv normal cortex regions (Fig. 5A, 64% and 47%, respectively, p < 0.001 for both). When the central ischemic core MAP2 IMR was compared to the MAP2 IMR in the dorsal and ventral ICp regions, statistically significant decreases of 72% (p < 0.001) and 74% (p < 0.001) were noted. However, there was no difference between the MAP2 IMR in K^+^-depleted and non-K^+^-depleted dorsal or ventral peripheral ischemic core regions (ICpd-DP vs. ICpd-ND: 0.464 ± 0.051, n = 11 vs. 0.478 ± 0.037, n = 8, p = 0.83; ICpv-DP vs. ICpv-ND: 0.493 ± 0.045, n = 10 vs. 0.419 ± 0.024, n = 9, p = 0.16, unpaired t-tests) in contrast to the differences observed in [K^+^]_br_ between non-K^+^-depleted and K^+^-depleted peripheral ischemic core regions. Thus, those regions with K^+^-depletion did not show a similar depression of MAP2 IMR.

## DISCUSSION

The main findings of this experimental study of brain potassium loss in ischemic stroke are: 1) that the loss of K^+^ from ischemic brain tissue agrees with previous measurements using less regionally specific methods; 2) that there were two statistically distinct populations of [K^+^]_br_ and K^+^-egress in the peripheral ischemic core regions; and 3) these differences in K^+^-dynamics are not related to pathological degradation of neuronal structure. These K^+^ changes in the peripheral regions of the ischemic core are at the crux of ischemic pathology with its hallmark of K^+^-mediated ischemic depolarization, which - with high [K^+^]_ex_ - was used to initially define the ischemic threshold^45^, the penumbra^46^, and ischemic SDs^46^ using K^+^-sensitive electrodes.

Our measurements of total brain K^+^, [K^+^]_br_, in ischemic tissue from histo-K^+^-stain imaging can be compared to [K^+^]_ex_ values obtained in a similar permanent MCAo model by Sick et al.^31^. However, Sick et al.’s^31^ measurements show a stability of [K^+^]_ex_ in ischemic core, while our and others’ measurements of [K^+^]_br_ show a linear decrease. To explore these different time courses we used the formalism of Cserr et al.^47^ to calculate [K^+^]_in_ from our [K^+^]_br_ values and the [K^+^]_ex_ values of Sick et al.^31^. The equations of Cserr et al.^47^ are based on the conservation of mass, fractional water content, and fractional extracellular space volume in depolarized brain^48,49^. Table 3 provides the results of these calculations using our estimates of central ischemic core [K^+^]_br_ at 0, 0.5, 1.5, and 3 h and [K^+^]_ex_ from central ischemic core Zones 2 and 3 and normal cortex Zone 8 (used as an estimate for t = 0) from Sick et al.^31^. The estimates of [K^+^]_in_ show a decrease over time and suggest that the fall of total brain [K^+^] was due to the decrease in [K^+^]_in_ until at 3 h [K^+^]_in_, [K^+^]_ex_, and [K^+^]_br_ are approximately equal. This equality indicated complete depolarization as confirmed by the estimated K^+^ Nernst potential of -5 mV. These estimates of [K^+^]_in_ are consistent with our observed decreases over time in [K^+^]_br_ in central ischemic cortex.

**Table 3.**
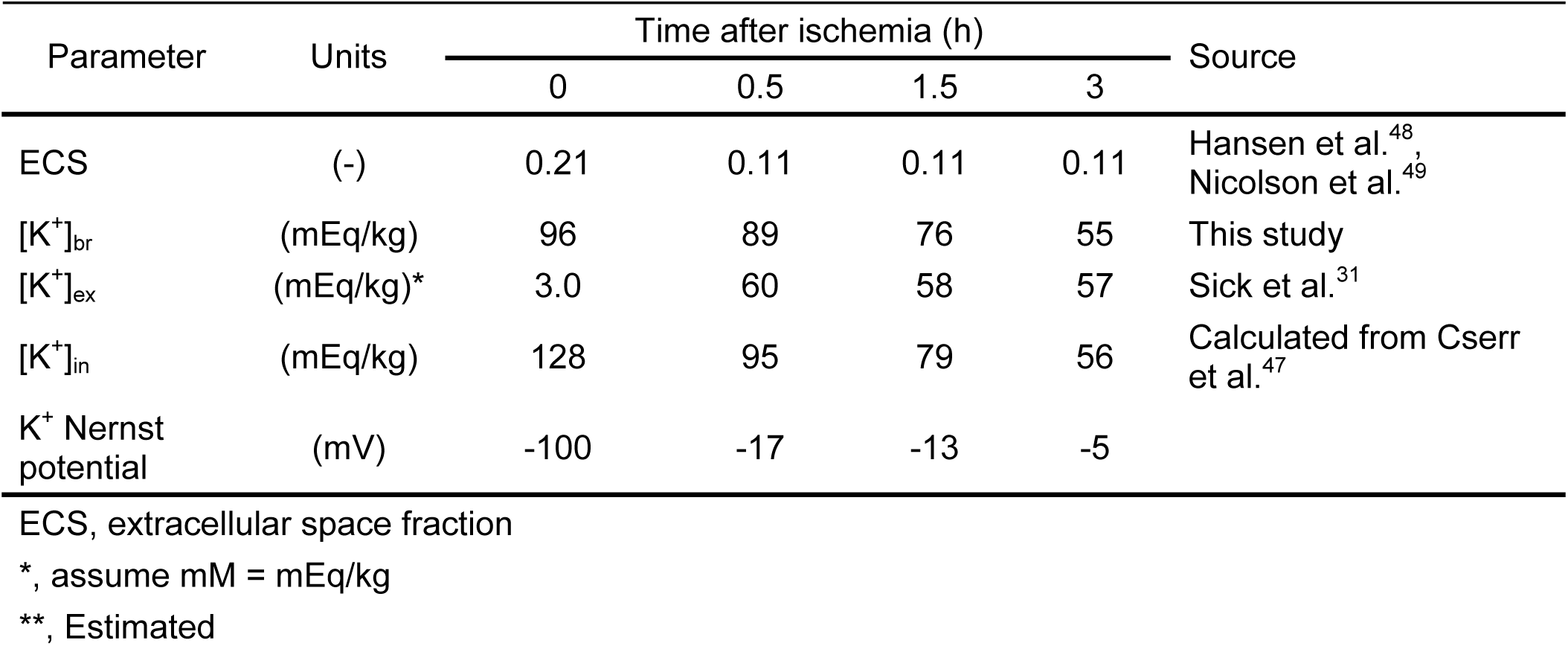
Estimates of [K^+^]_in_ in the central ischemic core.

| **Table 3** Estimates of $[K^+]_{in}$ in the central ischemic core | | | | | | |
| --- | --- | --- | --- | --- | --- | --- |
| Parameter | Units | Time after ischemia (h) |  |  |  | Source |
|  |  | 0 | 0.5 | 1.5 | 3 |  |
| ECS | (-) | 0.21 | 0.11 | 0.11 | 0.11 | Hansen et al.<sup>48</sup>,<br>Nicolson et al.<sup>49</sup> |
| $[K^+]_{br}$ | (mEq/kg) | 96 | 89 | 76 | 55 | This study |
| $[K^+]_{ex}$ | (mEq/kg)* | 3.0 | 60 | 58 | 57 | Sick et al.<sup>31</sup> |
| $[K^+]_{in}$ | (mEq/kg) | 128 | 95 | 79 | 56 | Calculated from Cserr<br>et al.<sup>47</sup> |
| $K^+$ Nernst<br>potential | (mV) | -100 | -17 | -13 | -5 | |

ECS, extracellular space fraction
\*, assume mM = mEq/kg
\*\*, Estimated

Our measurements of excessive loss of K^+^ from peripheral ischemic core regions suggest several possible consequences. The first is that isoelectric SDs could become more likely during this initial 5 h period after stroke onset as without K^+^, repolarization cannot occur. Alternatively, even if some regions that have undergone an SD near the ischemic core can repolarize, SD propagation into adjacent ischemic regions becomes improbable because K^+^ loss prevents the necessary repolarization. Even if the Na^+^,K^+^-ATPase could operate, there is not enough K^+^ left to re-supply [K^+^]_in_ and repolarize the cell membrane. This is consistent with Hartings et al.’s^35^ observations that SDs occur frequently (∼6/h) in the initial 2 h after stroke onset but then become less frequent (∼2/h) between 3-7 h. This suppression of SD occurrence after 2-3 h from stroke onset also suggests a temporary limit to SD-mediated infarct expansion^50^.

Perhaps the most important consequence of the loss of K^+^ near the ischemic penumbra involves the reduced chance of recovery. With such low [K^+^]_br_ there is not enough K^+^ left to reestablish [K^+^]_in_ even if circulatory supply is reinstituted and Na^+^,K^+^-ATPase functionality is re-established. Thus, the probable consequence of low K^+^ is that the membrane potential cannot be reestablished and neuronal/glial rescue is improbable: “Normalization of [K^+^]_e_ [after SD] is one of the critical components for restoration of neuronal activity”^51^. Young et al.^52^ suggested the direct replacement of K^+^ during neurosurgical procedures so that repolarization and neural transmission can occur although it is unclear how this could be accomplished. Two of these factors, limited infarct expansion and the decreased possibility of neuronal recovery, represent seemingly conflicting mechanisms that are in actuality a sequence of an immediate positive factor followed by a long-term negative. Thus, K^+^-depletion at the peripheral ischemic core has important clinical and therapeutic implications resulting from an enhanced understanding of the intra-ischemic core variations in the SD-mediated pathological progression of focal ischemia.

### Pathophysiology of the peripheral ischemic core

Our observations of regional variations in K^+^-egress in the ischemic core complement the changes in the peripheral ischemic core in the acute post-stroke time of up to 5 h as mentioned in the introduction^7,25–31^, but are not often the subject of studies of focal ischemic pathology and are distinct from the micro-heterogeneity and temporal progression in the ischemic core defined by molecular and metabolic studies and observations of vascular integrity^53^. However, K^+^ losses from the entire ischemic region during focal ischemia have been well documented using tissue analysis by atomic absorption spectrophotometry or flame photometry (see Table 1). The estimated rate of K^+^-egress of 12.2 mEq/kg/h for the entire ischemic cortex from the pooled ischemic core ROIs (ICc, ICpd, and ICpv) in this study is consistent with an overall average of others’ tissue sampling results of 10.1 mEq/kg/h (Table 1)^15–24^, providing a pathophysiological validation to our quantitative K^+^-imaging method. However, the very much smaller sample weights of our micro-punch methodology (0.545 mg), compared to the mean of the others from Table 1 (66 mg), made possible our intra-ischemic core analysis that disclosed a more nuanced view of K^+^-egress from the ischemic core.

The bimodal analysis separating the peripheral ischemic core regions with accelerated K^+^-egress, ICp-DP, from those with K^+^-egress slopes similar to those of ICc, ICp-ND, succeeded because it was based on the comparison of each peripheral region to its own central ischemic core region within the same cortical ribbon. The finding that the bimodal distribution was statistically significant permitted the identification of a breakpoint between the K^+^-depleted and non-K^+^-depleted peripheral ischemic core regions as the trough of the bimodal distribution at 36 mEq/kg (see Fig. 3). This distinction in turn allowed for the observations of the significantly different K^+^-egress rates for ICp-DP and similar rates for ICp-ND and ICc as depicted in Fig. 4. Over one-half (56%) of the peripheral ischemic core regions have lower [K^+^]_br_ and higher K^+^-egress than the remaining and at 2.5 h these higher K^+^-egress regions lost approximately twice as much K^+^ compared to those without K^+^-depletion (see Fig. 4).

Our quantitative imaging measurements confirm the qualitative observation of lower gray-values in histo-K^+^-stained sections in the peripheral, compared to the central, ischemic core, that was made in the gerbil model of focal ischemia^25^. [K^+^]_br_ values from cortical brain punch samples in this study^25^ did not reveal a significant decrease of [K^+^]_br_ in the peripheral ischemic core in comparison to the ICc when expressed as mEq/kg dry weight (p = 0.085). However, when converted to mEq/kg wet weight using their own regional brain tissue water contents, this difference between the two was significant (p = 0.0025). In another study^42^ a qualitative decrease in [K^+^]_br_ in the peripheral ischemic core is evident in their Fig. 2B in histo-K^+^-stained ischemic gerbil brain, but this K^+^-depletion was not mentioned or analyzed.

### Imaging vs. other methods

The quantitative imaging methodology used in this study has enabled a clear view of potassium dynamics in the ischemic core that has not been accessed by other methods including K^+^-sensitive microelectrodes, K^+^-sensitive fluorochrome imaging, and micropunch tissue sampling because it combines an appropriately high spatial resolution with a width breath of view. This view allowed for the definition of the border not only between the ischemic core and normal cortex but also between the peripheral and central ischemic core.

### Relation of K^+^-depletion to neuronal integrity

Although we observed depressed MAP2 IMR staining in the peripheral ischemic core, there was no exaggerated MAP2 depression in those peripheral ischemic core regions that were K^+^-depleted (ICp-DP) compared to peripheral core regions that were non-K^+^-depleted (ICp-ND). Both non-K^+^-depleted (ICp-ND) and K^+^-depleted regions (ICp-DP) had the same level of MAP2 IMR depression (see Fig. 5A). This lack of correspondence between the decrease in MAP2 IMR and K^+^-depletion is consistent with our contention that K^+^-depletion is not based on calpain-mediated microtubule disassembly and that the heterogeneity of K^+^-egress dynamics lacks a neuronally specific morphological correlate.

### Association of K^+^-depletion with SD occurrence

A ^42^K loading study showed that SDs cause massive amounts of K^+^ to leak from the brain to the surgically exposed brain surface^54^. This massive efflux of labeled K^+^ occurred in synchrony with the DC-shift of a remotely- and electrically-elicited SD, so is consistent with the K^+^-egress of our study because SDs occur with high frequency in the first hours after stroke^35^. The K^+^ efflux detected in Brinley et al.’s study^54^ was detected on the exposed brain surface, so does not address the K^+^-depletion we observed, although the high [K^+^]_ex_ from SDs is undoubtedly the first step in K^+^ leaving the brain.

### Possible routes of K^+^-egress

Once in the subarachnoid space CSF, K^+^ would have many possible egress routes. These egress routes include nerve sheaths, arachnoid villi/granulations, dural spaces adjacent to sinuses, and adventitial spaces of cranial foramina vessels^55^. One of the possible K^+^-egress routes from the subarachnoid space is via the meningeal lymphatics, as suggested by a study of VEGF-C overexpression^56,57^, and then via the cribriform plate into nasal lymphatics^58,59^ and superficial and deep cervical lymph nodes^56,60,61^. The K^+^-egress route to superficial, parotid, submandibular, and deep cervical lymph nodes would be similar to the egress pathway of lactate^62^.

The high [K^+^]_ex_ from SD occurrence in brain parenchyma would travel through the glia limitans superficialis into the subarachnoid space or would travel through the paravascular or glymphatic pathway^63,64^. Interstitial and CSF solutes, including K^+^, travel via this glymphatic pathway from the arteriolar perivascular space through astrocytes and out AQP4-containing astrocytic endfeet into venule perivascular spaces, eventually appearing in the subarachnoid space CSF or, alternatively, travel in the perivenular space to the nasal lymphatics. One feature of the glymphatic pathway that suggests K^+^ can move through the astrocytic endfeet is the colocalization of Kir4.1 ion channels, and perhaps other Kir channels, with AQP4 in orthogonal arrays of intramembranous particles^65,66^.

Although SD in normal brain has been shown to temporarily disable the glymphatic system flow as SD closes paravascular spaces^67^ due to astrocytic endfeet swelling^68^, the situation for ischemic SDs might well be different. Indeed, during the SDs of ischemia, the vasoconstriction known as cortical spreading ischemia that occurs as a result of high [K^+^]_ex_ and low NO^69^ provide a situation for increases in paravascular spaces^70^. This very early increase in perivascular space after ischemic onset^70^ occurs in the same time period during which K^+^-depletion occurs (before our 2.5 hour first data point in this study). The early influx of Na^+^ and water via the glymphatic system noted in this study^70^ speaks against this route also carrying the excessive increase in [K^+^]_ex_ out of the brain unless it’s bidirectional or the egress follows at a later time, but still within the 2.5 h that we show excessive K^+^ egress in Fig 4. Even though the glymphatic pathway is eventually compromised at 3 h after ischemic stroke^71^, activation of the superficial and deep cervical lymph nodes has been demonstrated at the end of this 3-h early acute period of experimental stroke in rats^72^.

We suggest that our use of 1% isoflurane plus 70% nitrous oxide did not interfere with the glymphatic system transport^73,74^ even though higher concentrations of 2.5% have been shown to effect this system^75^.

The slower decrease in K^+^-egress shown in Fig. 4 in the ICp-DP region after 2.5 h could be related to the dysfunction of the glymphatic system at this time even as other K^+^-egress systems are working. After MCA occlusion via ferric chloride initiated thrombosis, Gaberel et al.^71^ showed impaired glymphatic flow at 3 h. This study^71^ suggests that linear K^+^-egress after 2.5 h Is not associated with the glymphatic pathway.

The transcellular route through endothelial cells via abluminal^12,76^ and luminal^77^ K^+^ channels/transporters has been shown to be a possible K^+^ exit route, in addition to an Na^+^ and edema entrance route, within a 3 h time period after stroke onset^12^. This BBB-dependent route would lead to venous clearance of K^+^ from the peripheral ischemic core regions with more collateral flow potential. One other K^+^ exit possibility is not active in the initial time period of 5 h after stroke: The paracellular route via disrupted tight junctions in the endothelium^78^ for edema entry takes two days to become active^79^.

Varying degrees of K^+^-depletion from the peripheral ischemic core could suggest regional differences in K^+^-clearance that could be due to either more SDs and more [K^+^]_ex_ or differential access to K^+^-egress routes. Regional differences in [K^+^]_ex_ are certainly possible as watershed areas of marginal collateral circulation potential would have more SDs and higher [K^+^]_ex_ levels compared to regions with more vibrant collateral potential^80^. It might be that some of the non-K^+^-depleted regions are near the regions distal to the transected MCA, which might impair access to the initial glymphatic system pathway via the arterial paravascular space.

### Limited study of brain waste product egress

K^+^ has not been a subject of the many investigations of potential K^+^ egress routes extensively summarized by Rasmussen et al.^55^. “Only a limited number of endogenous brain waste solutes” (e.g., Ab^63^, tau^81^, and lactate^62,82^) “have been shown to clear via the glymphatic system”^83^. To our knowledge, the only specific mention of K^+^ clearance by the glymphatic system in the ischemic core is made in a figure caption of an editorial comment^84^ although the clearance of “solutes” (which of course include amyloid-beta as well as K^+^) has been widely discussed.

### Three-dimensional depiction of K^+^-depletion^34^

The histological section ribbon analysis decrements in [K^+^]_br_ as shown in Fig. 2E, 2H, and 5B have been previously presented three-dimensionally^34^. In the three-dimensional depiction of Fig. 6, these lower [K^+^]_br_ in the peripheral ischemic core form a torus-like region of low [K^+^]_br_ surrounding the ischemic core.

**Figure 6.**
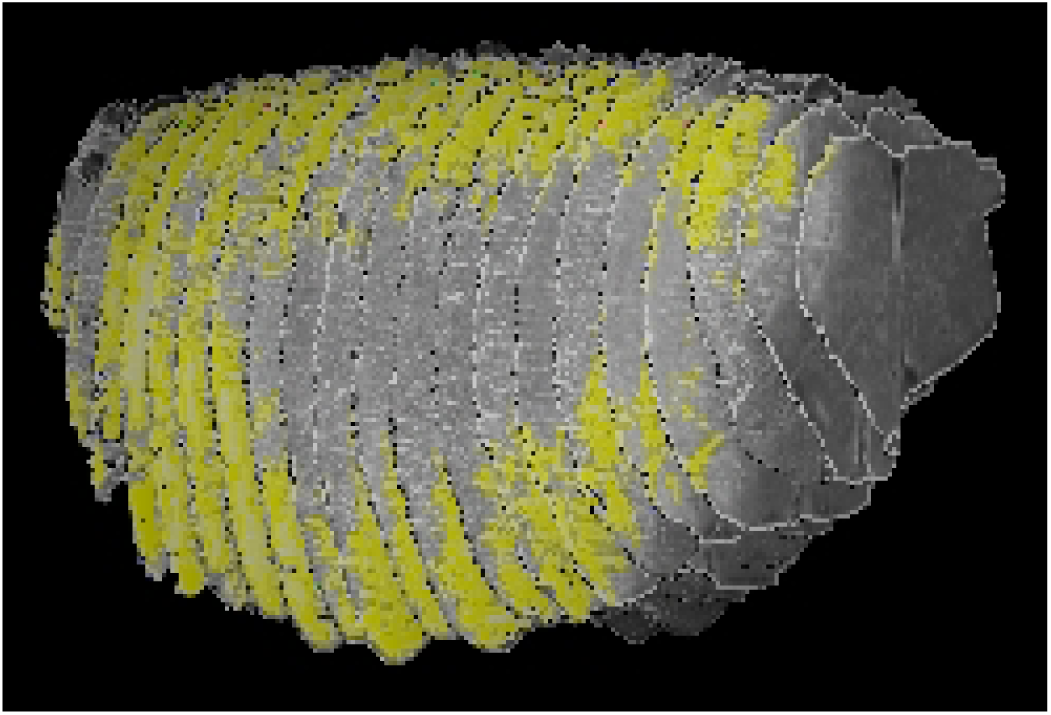
Three-dimensional reconstruction of a rat brain at 4.5 h after an intraluminal suture stroke from 21 stacked histo-K^+^-stained coronal slices (0.7-mm separation) as viewed from the front-right perspective. Regions with [K^+^]_br_ below 36 mEq/kg are shown in yellow color to highlight the peripheral ischemic core surrounding the central ischemic core. The rostral K^+^-loss in the peripheral ischemic core is less intense and narrower than at the caudal, ventral, and dorsal peripheral ischemic core. Permission granted from Yushmanov et al.^34^.

## Conclusion

Quantitative histo-K^+^ analysis after acute experimental focal ischemia revealed a greater decrease of potassium in a majority of peripheral ischemic core regions compared to the central ischemic core, possibly related to their proximity to K^+^-egress pathways. It should be clear that although we have demonstrated excessive regional variation in the well-known loss of K^+^ from the brain after ischemic stroke, it is unresolved as to what route, of the many possible, this K-egress takes. In addition, differences in these potential K^+^-egress pathways between rodent models and humans related to parasagittal dural lymphatics may exist^85^. These within-ischemic-core differences are important because the relatively greater K^+^-depletion from the peripheral ischemic core in comparison to the central ischemic core would temporally limit K^+^ availability for the further SD depolarizations that suggests a limit to SD-mediated secondary brain injury and would make neuronal recovery less likely even if reperfusion occurs.

## Supporting information

Supplemental Material - Table of Non-Standard Abbreviations

## Author contributions

AK developed the staining system and implemented the MCAt&biCCAo occlusion model. BY implemented the MCA-suture model. AK, VEY, and SCJ performed the experimental procedures. VEY wrote the first draft and contributed additional analysis. KAE conceived of and performed the statistical analysis. SCJ, AK, and VEY edited the manuscript. SCJ conceived of the project, provided supervision, analyzed data, prepared figures, and extensively revised the manuscript.

## Funding

This project was conducted at the Allegheny-Singer Research Institute, the previous work place of the authors. This work was partially supported by grants from the: US Public Health Service National Institutes of Health: NS30839, NS30839-14S1 and NS66292 to SCJ while at the Allegheny-Singer Research Institute; and Small Business Innovative Research grants 5R43NS092181 and 3R43NS092181-02S1 to SCJ for CerebroScope.

## Disclosure and Declaration of Competing Interests

Stephen C. Jones and Alexander Kharlamov are founding partners and shareholders of CerebroScope. However, nothing in this manuscript pertains to any commercial or financial activity of the company.

## Compliance with Ethical Standards

All ethical standards have been met.

## Data and Code Availability

Available upon request based on the submission and suitability of a formal project outline.

## Acknowledgements

We are grateful for the technical assistance provided by Jayjayantee Dasgupta and the careful reading of the manuscript by Thomas Ferguson.

## Supplemental Material

List of Abbreviations

## Notes

### Summary of Updates

Many details and minor errors have been corrected.

