## Supplemental Material - Table of Non-Standard Abbreviations for "Quantitative Imaging of the Heterogeneity of Brain Potassium Depletion in Experimental Focal Ischemia"

Table of contents:

List of Abbreviations ........................................................................................................................ 2

### Non-standard Abbreviations and Acronyms

$[K^+]_{br}$ = brain tissue potassium concentration (mEq/kg)

$[K^+]_{ex}$ = extracellular potassium concentration (mM)

$[K^+]_{in}$ = intracellular potassium concentration (mM)

$[K^+]$IC = $[K^+]_{br}$ of the ischemic cortex (mEq/kg)

$[K^+]$ICc = $[K^+]_{br}$ of the central ischemic cortex (mEq/kg)

$[K^+]$ICp = $[K^+]_{br}$ of the peripheral ischemic core (mEq/kg)

$[K^+]$ICp% = 100 x [$[K^+]$ICp / $[K^+]$ICc (%)

$[Na^+]_{br}$ = brain tissue sodium concentration (mEq/kg)

AIC = Akaike Information Criterion

BBB = Blood-Brain Barrier

CBF = Cerebral Blood Flow

CSF = CerebroSpinal Fluid

SD = cortical Spreading Depolarization

Gd-DTPA = gadolinium diethylenetriamine-pentaacetic acid

GV = gray value

Histo-$K^+$-stained = histochemically stained for $K^+$

ICc = central ischemic core

ICp = peripheral ischemic core

ICpd = dorsal peripheral ischemic core

ICpv = ventral peripheral ischemic core

ICp-DP = $K^+$-depleted peripheral ischemic core

ICp-ND = non-$K^+$-depleted peripheral ischemic core

IMR = immunoreactivity

LDF = laser Doppler flowmetry

MABP = mean arterial blood pressure

MAP2 = microtubule-associated protein 2

MCAt & biCCAo = middle cerebral artery transection and bilateral common carotid artery occlusion

NC = normal cortex

NCd = dorsal normal cortex

NCv = ventral normal cortex

PVS = perivascular spaces

ROI = regions-of-interest

Ta = time from stroke onset (h)

**Used in figures only:**

IH-ribbon = ischemic hemisphere cortical ribbon

NH-ribbon = non-ischemic hemisphere cortical ribbon

GLS – glia limitans superficialis

TD – terminal depolarization
